# Transient versican expression is required for β1-Integrin accumulation during podocyte layer morphogenesis in amphibian developing kidney

**DOI:** 10.1101/2025.07.20.665792

**Authors:** Isabelle Buisson, Jean-François Riou, Muriel Umbhauer, Ronan Le Bouffant, Valérie Bello

## Abstract

The functional organization of the vertebrate nephron is remarkably conserved. However, the morphogenetic processes underlying nephrogenesis can vary significantly across species and various kinds of kidneys. The *Xenopus* larval kidney, the pronephros, is a non-integrated nephron where plasma filtrates are first release into a coelomic compartment, the nephrocoel, before being released into the tubular compartment through ciliated funnels called nephrostome. Mechanisms of pronephros morphogenesis, especially the potential role of the extracellular matrix (ECM) remain poorly understood. This study investigates the role of the ECM component versican (vcan) in the development of the pronephric kidney in *X. laevis*, paying a special attention to non-integrated nephron features, the glomus, nephrocoel and nephrostomes. Vcan is dynamically expressed in the ECM surrounding the developing tubule and the podocyte layer of the glomus, with a transient presence in the differentiating podocyte-region prior to the formation of the concave podocyte pocket accumulating β1-integrin. Morpholino depletion of *vcan* causes a fusion of the proximal tubule branches without affecting the nephrostomes. Tubules become dilated and lose their proximal circumvolutions. More strikingly, *vcan* depletion results in a severe disruption of glomus morphogenesis, the podocyte layer failing to form its characteristic C-shaped structure. Despite normal expression of the differentiation markers *nphs2* in podocytes, β1-integrin fails to accumulate in the podocyte layer in vcan-depleted embryos. Interestingly, other ECM components, including fibrillin, laminin, and fibronectin, remain correctly localized, suggesting that the defect is not due to a general ECM disorganization. These findings reveal that transient vcan expression is critical for podocytes layer morphogenesis, likely by enabling β1-integrin accumulation and subsequent cell-ECM interactions necessary for structural assembly. The study highlights a specific and temporally regulated role of vcan in glomus morphogenesis, expanding our understanding of ECM dynamics in kidney development.

## Introduction

The kidney is a complex organ that performs essential functions such as blood filtration and fluid homeostasis. During development, three successive renal structures of increasing complexity arise from the intermediate mesoderm through inductive processes: the pronephros, the mesonephros, and metanephros, the latter being exclusive to mammals, birds, and reptiles [1]. Although kidney structures vary in their organization and complexity, they all share the nephron as the fundamental structural and functional unit. Notably, the functional segmentation of the nephron and gene regulatory network involved during kidney development are highly conserved among vertebrates [2][3]. Nephrons consist of a blood filtration unit (glomerulus or glomus when it spans several body segments) connected to a tubular compartment that enables ions and proteins reabsorption to form the definitive urine that is then excreted from the body. Two main types of nephrons have been identified, depending on whether the blood filtrate is released into the coelom or a compartment of the coelom (nephrocoel), or whether it goes directly into the tubule. The first type is called a non-integrated nephron and is characteristic of the pronephros of larval amphibians. The second type, the integrated nephron, is found in mesonephric and metanephric kidneys. It consists of a glomerulus enclosed within a Bowman’s capsule, which collects the filtrate from the blood before its passage into the renal tubule [4]. The pronephric kidney of *X. laevis* tadpoles is a well-characterized example of non-integrated nephron. It is a single nephron formed of a glomus spanning two or three body segments that protrudes into the nephrocoel where the plasma filtrate is collected. This filtrate is then redirected into the tubule through three ciliated funnels called nephrostomes. The definitive urine is eventually conducted through the duct to be released out *via* the cloaqua [5].

The filtration unit (glomerulus or glomus) is a complex structure composed of a fenestrated endothelium, a glomerular basement membrane (GBM), and podocytes with well-developed foot processes that form slit diaphragms [6]. The ultrastructure of this filtering barrier appears to be highly conserved among vertebrates, in both integrated and non-integrated nephrons. This has been demonstrated in the lamprey [7] as well as in teleost fishes such as medaka [8] and zebrafish [6]. It is therefore presumed to be conserved in the amphibian pronephros, although this has not been studied in detail. The gene network controlling glomerulus/glomus development is also largely conserved among vertebrates, with activation of *wt1* that regulates downstream genes such as *foxc2, hey1, mafb*, and *tcf21* to promote podocyte differentiation, followed by *lmx1b* contributing to structural maturation through the expression of *nephrin (nphs1)* and *podocin (nphs2)*. In contrast, the developmental sequences of glomerulus/glomus morphogenesis vary greatly across species [9]-[12], and whether the nephron is integrated or not. This sequence of events, largely documented in mammals, critically depends on interactions between the extracellular matrix (ECM) and cell surface receptors such as integrins [13] [14]. In mice, β1-integrin is essential for anchoring podocytes to the GBM through heterodimerization with α3-integrin (α3β1) [15] [16]. This integrin complex interacts with two major components of the GBM: laminin and collagen IV [17] [18]. However, in other species, the formation of the podocyte layer has been only scarcely described.

Versican (vcan), a hyaluronan (HA)-binding ECM component, is widely expressed in vertebrate connective tissues and interacts with HA, fibrillin, fibronectin, and β1-integrin [19]. While expressed at low levels in most adult human tissues (e.g., brain, blood vessels) [20], the *VCAN* gene is strongly upregulated during early development and pathological conditions [21]. Vertebrate orthologs play key roles in processes such as neural crest cell migration, cardiovascular development, skeletogenesis, and vasculogenesis [22]. In *X. laevis* embryos, vcan works as an inhibitor of neural crest cell migration and acts as a guiding cue by forming exclusionary boundaries to define neural crest cells migration pathways [23]. In mice, Vcan was shown to be essential for heart valve development and is later degraded from mature valve leaflets to ensure proper function [24]. Its cleavage by ADAMTS (a disintegrin and metalloproteinase with thrombospondin motifs) proteases generates bioactive fragments like versikine, which modulate ECM remodeling, signaling, migration, and apoptosis [25]. In zebrafish, Müller-Deile et al. (2016) demonstrated that knockdown of *vcan* by morpholino results in proteinuria, edema, podocyte effacement, and glomerular endothelial cell swelling with loss of fenestrations [26], but vcan does not appear to be essential for glomerular development, even though it plays a critical role for glomerulus maintenance.

In *X. laevis* pronephros, the podocytes are organized as a concave epithelial layer protruding into the nephrocoel [27] [5], which is revealed by differentiation markers such as *nphs*2, *ptpru, nphs1* and *kirrel* [9]. Yet, morphogenesis of the glomus and the nephrocoel has not been described in detail. Notably, the potential role(s) of ECM components remain unexplored. In this study, we analyze the role of vcan in the development of the pronephric nephron, with a focus on the glomus and nephrocoel region. Expression studies reveal that vcan expression in the ECM is dynamic during the development of the pronephros, vcan being detected at early stages of glomus development in the ECM lining the developing podocytes and later disappearing when they display a high expression of ß1-integrin. Using a loss-of-function approach, we further show that *vcan* knockdown results in an abnormal development of the convoluted tubule leading to the formation of an enlarged tubule at tadpole stage, surrounded by correctly organized ECM containing fibrillin and laminin. The glomus is severely affected, with the podocytes layer failing to organize as a ß1-integrin expressing C-shaped pocket. These later results suggest that a tightly regulated spatio-temporal expression of vcan is essential for β1-integrin accumulation and proper assembly of the podocytes layer.

## Materials and Methods

### Antibodies

The vcan antibody has been obtained from Eurogentec. Briefly, a peptide corresponding to the KHHHSWSRTWQDSPR sequence, located in the C-terminal domain of vcan, was injected into guinea pig. The final bleed serum was purified by HPLC to obtain a purified antibody according to Eurogentec protocol. Commercial antibodies references are given in table 1.

**Table 1:** List of antibodies.

| Epitope | species | dilution | company |
| --- | --- | --- | --- |
| β-integrin | mouse | 1/50 | (8C8) DSHB |
| Fibronectin | mouse | 1/100 | (4H2) DSHB |
| Fibrillin | mouse | 1/50 | JB3 (DSHB) |
| Laminin | rabbit | 1/100 | Sigma L-9393 |
| XVersican | guinea pig | 1/50 | Eurogentec (homemade) |
| 3G8 | mouse | 1/40 | Gift from E. Jones |
| Alexa Fluor™ 488/568 goat anti-Rabbit IgG (H+L) | goat | 1/400 | A11008/A11036 ThermoFisher |
| Alexa Fluor™ 488/568 goat anti-Mouse IgG (H+L) | goat | 1/400 | A11001/A11031 ThermoFisher |
| Alexa Fluor™ 488/568 goat anti-Guinea Pig IgG (H+L) | goat | 1/400 | A11073/A11075 ThermoFisher |
| Anti-Digoxigenine AP fab-fragments | sheep | 1/2000 | A11093274910 ThermoFisher |

### Xenopus embryos, microinjection

*X. laevis* were purchased from TEFOR Paris-Saclay (France). Embryos were obtained after artificial fertilization and were raised in modified Barth’s solution (MBS). Stages were determined according to the normal table of *X. laevis* [28]. All animal experiments were carried out according to approved guidelines validated by the Ethics Committee on Animal Experiments “Charles Darwin” (C2EA-05) with the “Autorisation de projet” number 02164.02.

Microinjections were performed at the 4-cell stage as described previously [29], with vcan-Mo1 (5’-GTACAGGAATCTCCCCCAGACTCGT-3’), vcan-Mo2 (5’-TCTTGACCTTTTAAGGTGACCTAGT-3’) or the standard control morpholino (5’-CCTCTTACCTCAGTTACAATTTATA-3’) (Gene Tools). Morpholinos were injected into both left blastomeres at the 4-cell stages (15 ng/blastomere for vcan-Mo1 and standard control Mo and 50 ng/blastomere for vcan-Mo2).

### *In situ* hybridization and embryo sectioning

Whole-mount *in situ* hybridization (WISH) was carried out as previously described [30]. To generate plasmids used for antisense cRNA probe synthesis, probe DNA was generated from a stage 32 cDNA library using specific primers for *vcan* (forward 5’-ACAAATGAGTTCCTGCGAATCA-3’; reverse 5’-ACTCGGGTTGTGTGCTTGAAG-3’) and for *nphs2* (forward 5’-CCACAGAAACAGAGGGAACACA-3’; reverse 5’-TTGATCTCAGGCAGCCTGGT-3’). PCR products were cloned into the pCRII vector according to the manufacturer instructions (Invitrogen), and digoxigenin-labelled cRNA probes were transcribed from linearized plasmids using SP6 or T7 RNA polymerase. Embryos selected after WISH were sectioned as described in Bello *et al*. (2008) [31]. Briefly, embryos were washed three times in PBS, embedded in PBS containing 15% fish gelatin 15% sucrose, and frozen by immersion in isopentane. Sixteen μm thick frozen sections were prepared using a Leica CM3050S cryostat. Observation and image acquisition were performed on a Zeiss MacroApotome (Axiozoom V16).

### Immunodetection

Whole-mount immunochemistry with the 3G8 antibodies (kindly provided by Dr. E.A. Jones, Warwick University, United Kingdom) was performed using standard methods on MEMFA-fixed embryos. The secondary antibodies were alkaline phosphatase-conjugated goat anti-mouse. 5-Bromo-4-Chloro-3-Indolyl Phosphate/nitroblue Tetrazolium (BCIP*/*NBT) solution was used for the color reaction according to manufacturer’s recommendations (ROCHE 11681451001). Embryos selected after immunochemistry were sectioned as described in Bello *et al*. (2008) [31]. Sixteen μm thick frozen sections were obtained using a Leica CM3050S cryostat. Imaging was performed using a Zeiss MacroApotome (Axiozoom V16).

Immunofluorescence on embryos transverse sections were performed as previously described in Bello et al. (2008) [31]. Briefly, embryos were fixed in MEMFA solution for 1 hour, washed three times in PBS, and embedded in gelatin for cryostat sectioning. Fourteen μm cryostat sections were made and processed for immunodetection. Primary antibody concentrations are indicated in Table 1. Dilutions of secondary antibodies are indicated in table 1. For vcan immunostaining, an additional antigen retrieval step was included, involving boiling the sections for 20 minutes in 10 mM citrate buffer (pH 6.0). Each phenotype shown in the figures is representative of observations from an average of 5–8 embryos sectioned.

Fluorescence image were acquired using a Zeiss Observer Z1 microscope equipped with an Axiocam 506 monochrome cooled CCD camera, a 20x objective with (NA 0.8), and a 40x oil immersion objective (NA 1.3). All images were processed using Zen software (Blue Edition). Image analyses and figure preparation were carried out using Adobe Photoshop version 24.0.0.

### Statistical analysis

For figure 2, the *Nphs2*-stained area was measured using Fiji software to calculate the surface ratio between the injected side and the uninjected side of every analyzed embryo. Graphical dispersion and analysis were performed using R Commander (R software). A non-parametric Mann-Whitney test was used for significance.

## Results

### Expression pattern of vcan during pronephros development

The expression profile of *vcan* mRNA was previously described by Casini et al., 2018 [32]. As a first step, we aimed to further characterize in detail *vcan* mRNA and protein expression during *X. laevis* pronephros morphogenesis. This was carried out at the tailbud and tadpole stages using homemade vcan *in situ* hybridization probe and antibody (see material and methods and Fig 1).

**Fig 1.**
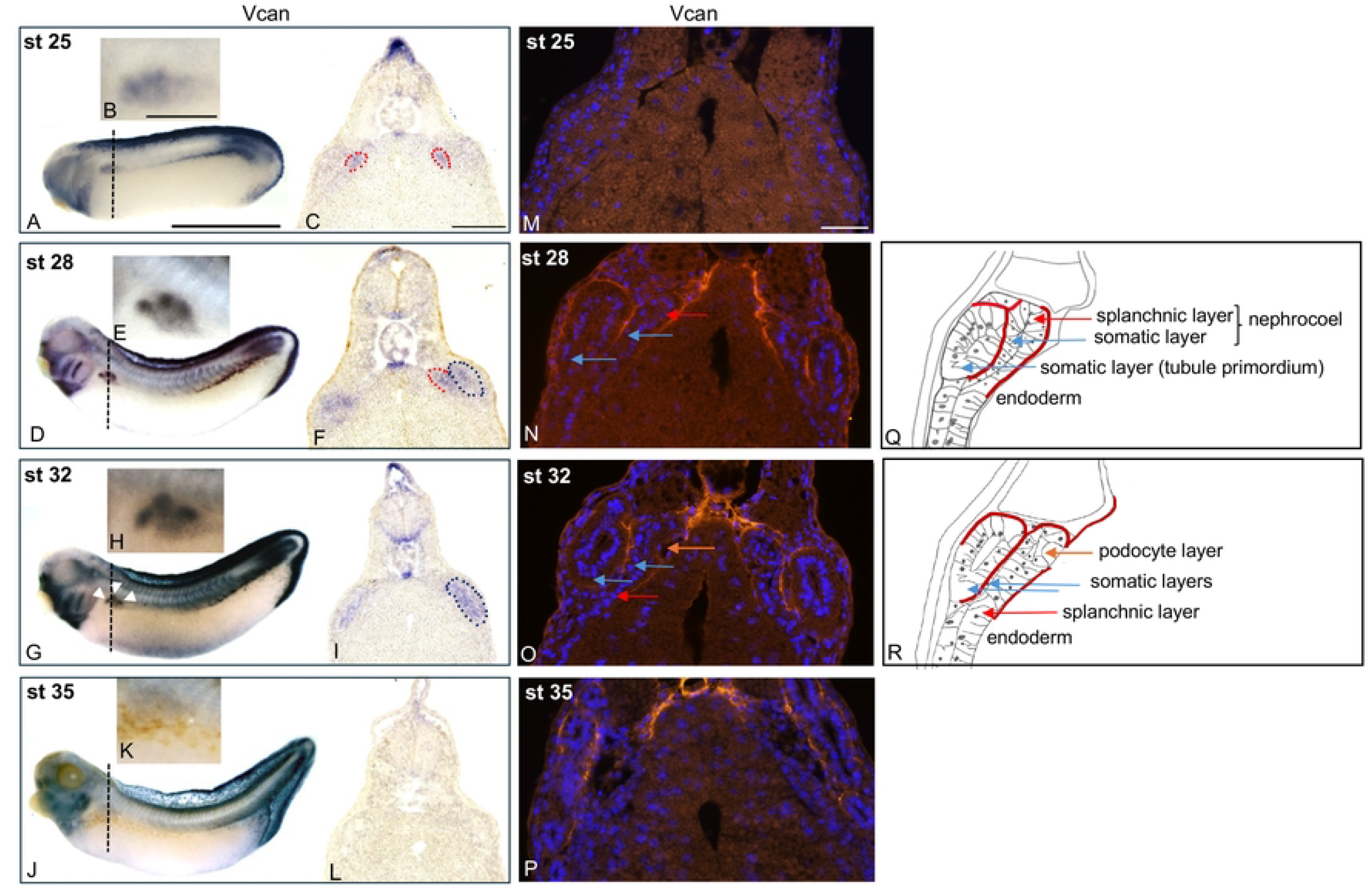
Spatial and temporal expression of vcan mRNA and protein during pronephric morphogenesis. (A-L) *Vcan* expression in *X. laevis* embryos by whole-mount *in situ* hybridization (WISH) from stage 25 to 35. The embryos are shown in lateral view (A, D, G, J) with a magnification of the pronephric region (B, E, H, K). Transverse sections made at the level of the pronephric region (black dashed line) are shown (C, F, I, L) (note that the sections were not made from the whole mount specimen shown). The glomus and pronephric tubules areas are delineated by red and blue dashed lines, respectively. Scale bars: 1.7 mm for whole embryo, 0.3 mm for embryo magnification and 280 μm for section. (M-R) Immunodetection of vcan protein in transverse sections from stage 25 to 35 (M, N, O, P). Nuclei are stained with DAPI (blue). Scale bar: 280 μm. (Q, R) schematic representation of transverse sections shown in N, O. Blue and red arrows in N and Q, and in O and R respectively, indicate the somatic and splanchnic layers. The orange arrow in O and R points to the podocytes layer, highlighting its concave appearance.

Low levels of *vcan* mRNA start to be detectable at stage 25 in the splanchnic layer which will participate to the podocyte layer of the glomus (Fig 1A, 1B and 1C). At this stage, the protein is not detected (Fig 1 M). From stage 28 onward, *vcan* mRNA expression strongly increases in the proximal region of the forming tubule (Fig 1D and 1E) and persists in the glomus area (Fig 1F). At stage 28, the pronephric anlage consists of the tubule primordium developing from the outer somatic mesodermal layer, which faces the epidermis, while the podocyte layer of the glomus emerges from the splanchnic mesodermal layer adjacent to the endoderm (Fig 1Q). The nephrocoel cavity which connects the tubule to the glomus will later appear between the somatic and splanchnic layers [5] [33]. The vcan protein is detected in the dorsal and lateral part of the ECM surrounding the developing tubule, especially at the contact of the somatic mesodermal layer that will contribute to the future wall of the nephrocoel. Vcan is also localized between the splanchnic mesoderm and the endoderm (Fig 1N and 1Q). At stage 32, a region of the dorsal splanchnic layer invaginates to form a concave structure where podocytes will further contact the vascular component of the glomus (Fig 1R). The vcan protein is clearly detected between the forming tubule and the somatic layer lining the future nephrocoel. It is also present at the interface between the splanchnic layer and the endoderm, except at the level of the concave structure corresponding to the podocyte layer (Fig. 1O and 1R). *Vcan* mRNA remains strongly expressed in the proximal part of the tubule but is no longer detected in the glomus area (Fig 1G, 1H and 1I). By stage 35, *vcan* transcripts are no longer detected in the pronephric region (Fig 1J, 1K and 1L), while vcan protein remains visible surrounding the proximal tubules (Fig 1P).

These observations reveal a dynamic expression of vcan transcripts and protein at key stages of pronephric kidney development, pointing to a role of vcan in the morphogenesis of this organ.

### Vcan depletion impairs tubule and glomus morphogenesis

To study vcan function in kidney development, we performed loss-of-function experiments by injecting into the two left blastomeres of 4-cell stage embryos various translational blocking morpholino targeting *vcan*. We have tested a morpholino previously described (vcan-Mo2) [23] as well as a newly designed morpholino (vcan-Mo1). Injection of vcan-Mo1 and vcan-Mo2 both resulted in a loss of vcan protein expression (S1 Fig). However, while vcan-Mo1 was effective at a standard concentration (30 ng per embryo), vcan-Mo2 required a higher dose (100 ng per embryo), as previously reported [23], to achieve a similar effect (S1 Fig). This high dose resulted in significant embryonic lethality. Therefore, we have chosen to use vcan-Mo1 for the functional investigation of vcan.

To assess pronephric kidney defects associated with *vcan* loss-of-function, we first analyzed the effect resulting from vcan depletion on proximal tubule differentiation at the early tadpole stage 40 using a 3G8 antibody staining [33]. In st-Mo-injected control embryos, no difference was observed between uninjected and injected sides (92%, n=50), with 3G8 staining revealing a highly convoluted proximal tubule (Fig 2A, 2B, 2C and 2D). In contrast, 3G8 staining revealed a differentiated tubule with a reduced proximal circumvolution on the vcan-Mo1 injected side (80%, n=46) compared to the uninjected side of the same embryo (Fig 2E and 2F). This effect was confirmed on transverse sections that further revealed a dilated proximal tubular structure (Fig 2G and 2H) on the injected side.

**Fig 2.**
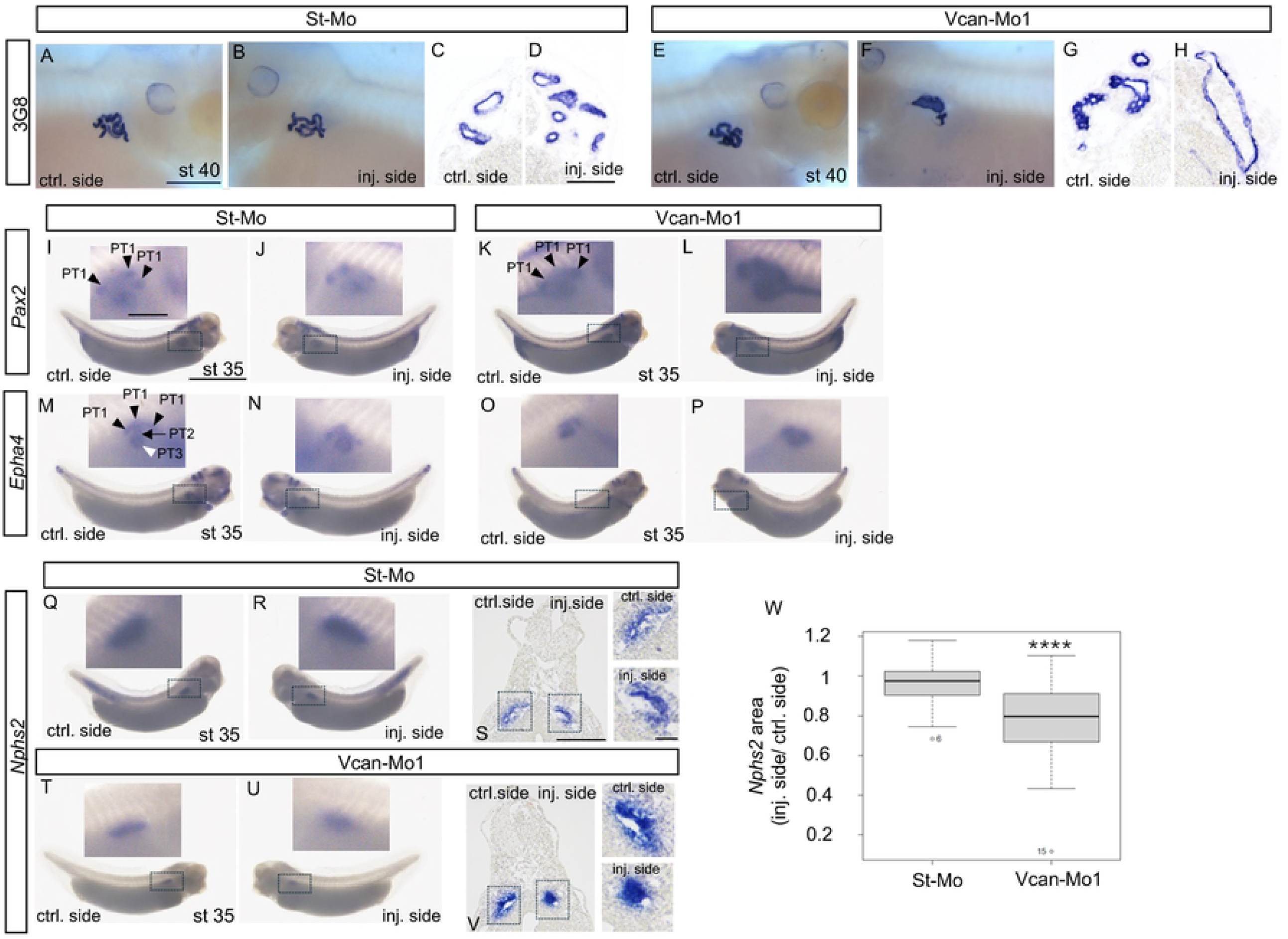
Vcan is required for glomus and tubules development. (A-H) 3G8 immunostaining at stage 40. (A, B) St-Mo injected embryo; (C, D) transverse sections of a st-Mo injected embryo. (E, F) Vcan-Mo1 injected embryo; (G, H) transverse sections of a vcan-Mo1 injected embryo. Scale bars: 280 μm for whole embryo and 93 μm for section. (I-L) WISH for the tubule marker *pax2* at stage 35. (I, J) St-Mo injected embryo; (K, L) vcan-Mo1 injected embryo, black arrows point to PT1 segments. (M–P) WISH for the tubule marker *epha4* at stage 35. (M, N) St-Mo injected embryo; PT1, PT2 and PT3 segments are indicated by black arrowheads, black arrow and white arrowhead, respectively. (O, P) Vcan-Mo1 injected embryo. (Q-W) WISH for glomus marker *nphs2* at stage 35. (Q, R) St-Mo injected embryo; (T, U) Vcan-Mo1 injected embryo; (S, V) transverse sections of st-Mo and vcan-Mo1 injected embryos. Scale bars: 1.7 mm for whole embryo, 0.3 mm for embryo magnification, 280 μm for section and 93 μm for section magnification. For all images: representative embryos are shown. Sections were not made from the whole-mount specimen shown. Magnification of the pronephric region is highlighted by the dotted squares. (W) Quantification of the nphs2-positive area (dotted regions in Q, R and T, U). The graphical dispersion displays the ratio between the injected and uninjected sides for each embryo (n=80). Statistical significance was assessed using the non-parametric Kruskal - Wallis test (**** indicates p < 0.0001).

Next, we investigated tubulogenesis defects in more detail by analyzing the expression of specific tubular segment markers, *pax2* and *epha4* at stage 35. *Pax2* expression was unaffected in st-Mo injected embryos (Fig 2I and 2J), with a proper arrangement of the three PT1 segments (90%, n=47) whereas comparison between the injected and uninjected sides revealed that *vcan* knockdown strongly affected the *pax2* expression domain (Fig 2K and 2L) (63%, n=48). On the control side, *pax2* was strongly expressed in all three nephrostomes and at lower levels along the rest of the proximal tubule. On the injected side, *pax2* expression in the nephrostomes was not clearly distinguishable from the rest of the proximal tubule, due to a stronger and more diffuse *pax2* signal in the whole proximal tubule (Fig 2L). This did not result from an alteration of the nephrostomes which normally expressed the *odf3* differentiation marker (S2 Fig). St-Mo injected embryos displayed normal *epha4* expression, showing proper organization of each segment of the proximal tubule PT1, PT2 and PT3 (Fig 2M and 2N) (85%, n=77), similar to the uninjected side of vcan-depleted embryos (Fig 2O). In contrast, on the vcan-depleted side, *epha4* expression revealed that the proximal segments are not properly individualized (Fig 2P) (84%, n=60).

To assess the role of *vcan* during glomus development, we analyzed the effect of vcan depletion on the expression of *nphs2*, which encodes the podocyte terminal differentiation marker podocin. As shown in Fig 2T and 2U, *nphs2* labeling revealed that 70% (n=47) of embryos injected with vcan-Mo1 exhibited a reduced expression domain compared to the control side. In contrast, only 10% (n=58) of embryos injected with the standard control morpholino showed a similar reduction (Fig 2Q and 2R). The ratio of injected to uninjected *nphs2* labelling area in the glomus region showed a significant reduction in vcan-Mo1 injected embryos (n=35) compared to standard control morpholino-injected embryos (n=45) (Fig 2W). To further investigate the effect of vcan depletion on glomus development, transverse sections of representative embryos were analyzed. On the control side, the podocyte layer expressing *nphs2* was organized into the characteristic concave structure (Fig 2S). On the vcan-Mo1 injected side, this structure appeared strongly reduced (Fig 2V). *Nphs2*-expressing cells were clustered near a small cavity. This phenotype was consistently observed across multiple sections from 7 different embryos analyzed (80%). Similar results were obtained following vcan-Mo2 injection (S3 Fig). The injection of vcan-Mo2 led to a reduction in the *nphs2* expression domain (53%, N=57), compared to the st-Mo-injected side (9%, N=64) (S3 Fig). These findings suggest that vcan depletion does not impair podocyte differentiation but rather disrupts the proper spatial organization of the podocyte layer.

In conclusion, vcan loss-of-function impairs both proximal tubule morphogenesis and the organization of the podocyte layer.

### Dynamic distribution of vcan ECM partners during early pronephric development

Through its interaction with various ligands, vcan plays a key role in ECM organization and in dynamic cellular processes during tissue and organ morphogenesis. To better understand the role of vcan in glomus and tubule morphogenesis, we analyzed the spatiotemporal expression of the major vcan partners β1-integrin and fibrillin during glomus and pronephric tubule development. To this end, co-immunostaining of vcan with either β1-integrin or fibrillin was performed on transverse sections of stage 28 and stage 32 embryos (Fig 3).

**Fig 3.**
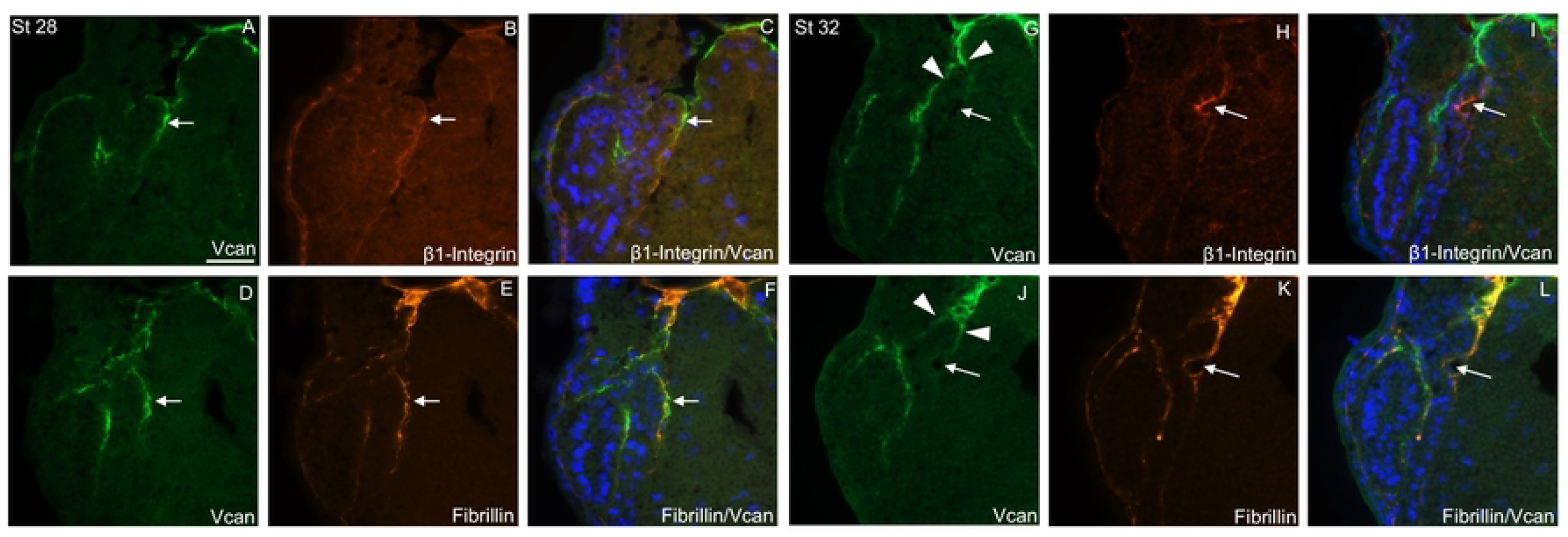
Expression of β1-Integrin, fibrillin and vcan in pronephric tubule and glomus region at stage 28 and 32. (A-F) Transverse sections of stage 28 embryo. (A-C) Co-immunostaining of vcan (A) and β1-integrin (B) and merged images with DAPI (C). (D-F) Co-immunodetection of vcan (D) and fibrillin (E) and merged images with DAPI (F). White arrows indicate the splanchnic layer, which will give rise to the podocytes layer. (G-L) Transverse sections of stage 32 embryo. (G-I) Co-immunodetection of vcan (G) and β1-integrin (H) and merged images with DAPI (I). (J-L) Co-immunodetection of Vcan (J) and fibrillin (K) and merged images with DAPI (L). White arrows point to the podocyte layer. White arrowheads point to the dorsal part of the glomus. Scale bar: 93 μm.

At stage 28, β1-integrin was strongly expressed basolaterally in the epidermal cells of the sensorial layer, as well as at the level of the pronephric splanchnic layer adjacent to the endoderm at the cell basal plasma membrane. It is also detected at the borders of other cell types, but at much lower levels (Fig 3B). Fibrillin and vcan expression patterns largely overlapped (Fig 3D, 3E and 3F).

Fibrillin was detected between the tubule anlage and the somatic layer that will contribute to the future wall of the nephrocoel, as well as in the dorsal region of the forming tubule. Fibrillin was also observed at the interface between the splanchnic and endodermal layers (Fig 3E). Interestingly, vcan, β1-integrin, and fibrillin showed partial co-localization in the region that will give rise to the podocyte layer (Fig 3C and 3F, white arrow).

At stage 32, β1-integrin expression was prominent at the site of the concave structure emerging in the pronephric splanchnic layer where podocytes are differentiating (Fig 3H). It is located at the cell surface facing the cavity that will be colonized by capillaries. Interestingly, this expression can be related to the established role of β1-integrin as a major ECM receptor essential for mammalian glomerular development and maintenance of structural integrity [13]. Fibrillin was detected around the pronephric tubule as well as between the splanchnic layer and the endoderm (Fig 3K). In contrast to vcan, fibrillin was detected in the ECM lining the cavity, which also expressed fibronectin and laminin (Fig 3J, 3K and 3L) and (S4 Fig).

These observations reveal that prior to the invagination of the podocyte layer, vcan, fibrillin and β1-integrin are all expressed along the splanchnic layer. Later, vcan expression disappears from the developing podocyte layer, where β1-integrin shows strong accumulation.

### Vcan is required for podocyte layer morphogenesis

To investigate the role of vcan in the assembly of fibrillin-containing ECM and β1-integrin expression during glomus and proximal tubule formation, immunostaining for β1-integrin and fibrillin was performed on vcan depleted embryos at stage 32 (Fig 4). Immunostaining of β1-integrin on transverse sections of st-Mo injected embryos revealed strong accumulation of β1-integrin in the invaginating podocyte layer on the injected side (Fig 4C), similar to uninjected control side (Fig 4B) and to control embryos described above (Fig. 3H). On the uninjected side of vcan depleted embryos, strong vcan expression was observed between the proximal tubule and the wall of the coelomic cavity (Fig 4G). Vcan was also detected in the dorsal part of the glomus region (Fig. 4G), without overlapping with β1-integrin which distinctly marked the concave podocyte layer (Fig 4E and 4H, white arrows). In contrast, on the injected side of vcan depleted embryos, β1-integrin failed to accumulate at the level of a concave structure, which did not form (Fig 4F and 4I, red arrow). Nevertheless, fibrillin is detected at the level of a concave structure in st-Mo (Fig 4J, 4K and 4L) and on the uninjected side (Fig 4P and 4Q), and remained detectable on the injected side of vcan depleted embryos where the missing concave podocyte layer normally takes place (Fig 4R).

**Fig 4.**
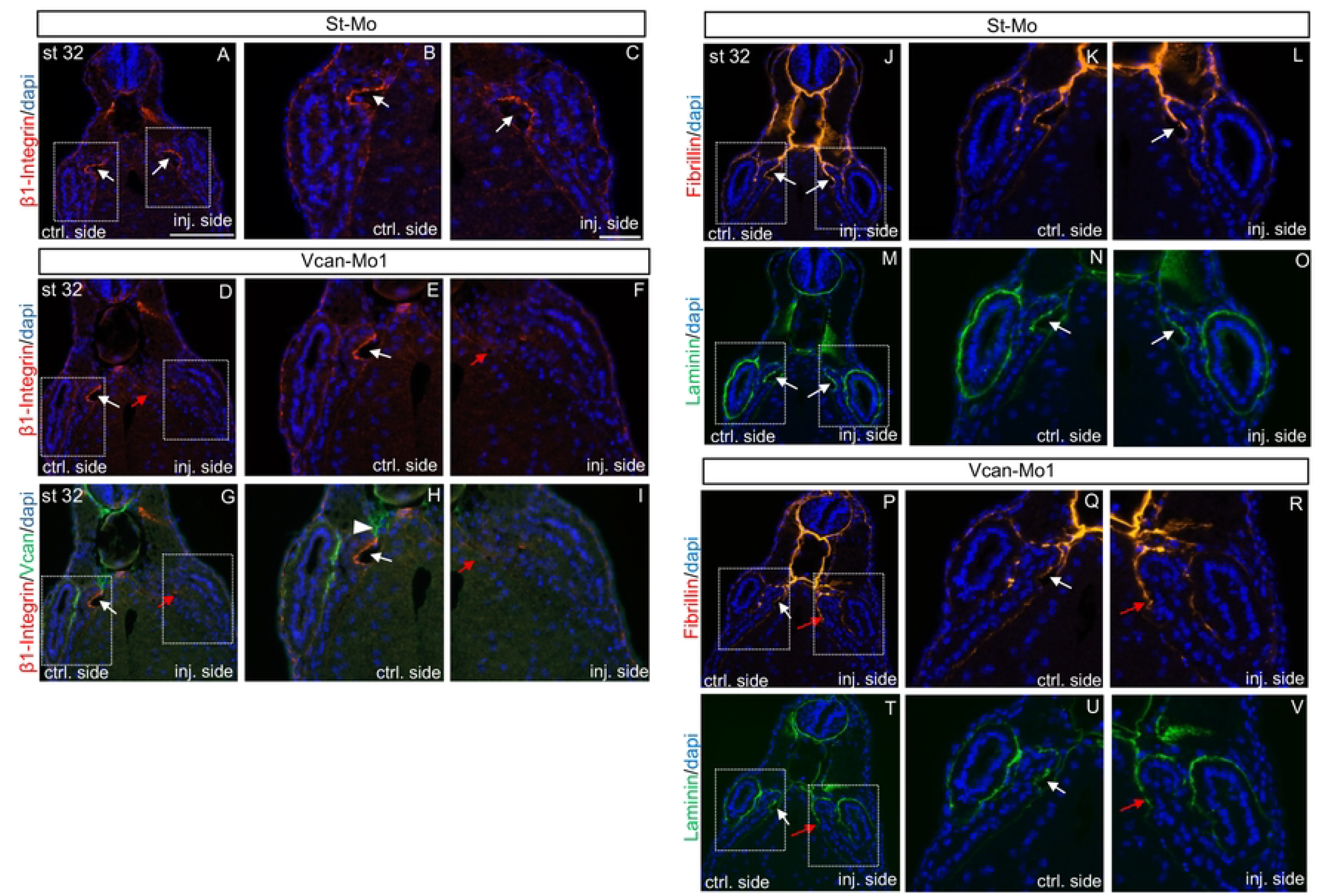
Vcan loss-of-function affects podocytes layer morphogenesis. (A-C) Transverse sections of st-Mo-injected embryo with β1-integrin immunostaining at stage 32. Nuclei are stained with DAPI (blue). (D-I) Co-immunostaining of β1-integrin and vcan on transverse sections of a vcan-Mo1 injected embryo at stage 32. (D-F) β1-integrin and DAPI. (G-I) Merged images with β1-integrin, vcan and DAPI. Panels (B, C), (E, F) and (H, I) are magnification of the pronephric areas indicated by dotted squares in (A), (D) and (G), respectively. White arrows indicate the localization of the podocyte layer. Red arrows point to the defects in the podocyte layer in panels (D, F) and (G, I). (J-O) Transverse sections of a st-Mo injected embryo with fibrillin (J-L) and laminin (M-O) immunostaining at stage 32. (P-V) Transverse sections of a vcan-Mo1 injected embryo with fibrillin (P-R) and laminin (T-V) immunostaining at stage 32. Nuclei are stained with DAPI (blue). White arrows point to the podocytes layer; Red arrows indicate podocyte layer defects. Panels (K, L), (N, O), (Q, R) and (U, V) are magnification of the pronephric area indicated by the dotted squares in (J), (M), (P) and (T), respectively. Scale bars: 280 μm for section and 93 μm for sections magnification.

The effect of vcan depletion on ECM proteins laminin and fibronectin, which are two key ECM components involved in early embryonic basement membrane assembly and cell differentiation, was further investigated. Similar to fibrillin, laminin was detected in the concave structure formed by the podocyte layer as well as around the proximal tubule on both the uninjected and injected sides of st-Mo injected embryos (Fig 4N and 4O). On the injected side of vcan depleted embryos (Fig 4V), laminin expression persisted at the site where the podocyte layer would normally form. A similar distribution was observed for fibronectin (S4 Fig).

Together, these results indicate that in the absence of vcan, the podocyte layer fails to organize properly into the characteristic pocket protruding into the nephrocoel. This defect may be related to the lack of β1-integrin accumulation in this region. Yet, the localization of other ECM components such as fibrillin, laminin and fibronectin appears unaffected, indicating that defects in the organization of the podocyte layer does not simply result from a gross absence of the ECM lining the splanchnic layer. Similarly, the ECM surrounding the proximal tubule also appears to remain intact.

## DISCUSSION

In this study, we demonstrate that vcan plays a key role in *Xenopus* pronephric kidney development. Vcan is dynamically expressed in the ECM surrounding the developing tubule and podocyte layer, with a transient enrichment in the region forming the concave podocyte pocket. Loss of vcan function affects the individualization of the proximal tubule segments and severely impairs the organization of the glomus podocyte layer, as evidence by the absence of a pocket accumulating ß1-integrin, despite the continued expression of the podocyte terminal differentiation marker *nphs2*. Our findings suggesting that vcan is essential for proper glomus morphogenesis in the developing *Xenopus* pronephros contrast with those reported for the zebrafish pronephros where vcan depletion was not found to affect glomerulus development per se, but rather the maintenance of glomerulus function at later stages [26]. This is likely related to the different processes taking place during zebrafish glomerulus and *Xenopus* glomus development. During development of the zebrafish pronephros, podocyte precursors migrate toward the midline and recruit endothelial cells to form a single vascularized glomerulus shared by two nephrons [8]. In *Xenopus*, a podocyte layer rather protrudes inside the nephrocoel and interacts with the vascular components evaginating from the dorsal aorta [4] [5]. The different function of vcan observed in our study may therefore reflect distinct requirement of vcan in the establishment of the filtration barrier in these two species. The phenotype we describe may be specific to the non-integrated glomus structure characteristic of the *X. laevis* pronephros.

At stages 28 and 32, vcan, fibrillin and laminin show partial colocalization in the ECM which separates the tubule from the coelomic wall within the somatic layer, as well as in the splanchnic layer. Fibronectin was also detected, exhibiting a broader expression domain than either fibrillin, laminin or vcan (S4 Fig). Following vcan depletion, the distribution of fibrillin, fibronectin and laminin, remained unchanged, suggesting that vcan is not directly required for ECM organization in the developing glomus. These observations rather suggest that fibrillin, fibronectin and laminin containing ECM are organized in the developing pronephros first before vcan deposition. This hypothesis is consistent with the literature indicating that fibronectin, fibrillin, and laminin assemble sequentially and in an interdependent manner within the extracellular matrix. Vcan, in contrast, is incorporated at a later stage by interacting with already organized networks, primarily through fibrillin [19] [34] [35].

Our results reveal a tightly regulated spatio-temporal expression of vcan during podocyte layer formation. At early stages of glomus development (stage 28), vcan localizes to the region where the podocyte layer will form, but its expression disappears during podocyte layer assembly by stage 32. However, what remains unclear is how the disappearance of vcan is mechanistically linked to the podocyte layer formation at stage 32, when this structure adopts its characteristic concave morphology.

A hypothesis to link the disappearance of vcan to the formation of the concave podocyte region relies on its role in modulating ECM properties. Previous studies have shown that a tightly regulated temporal and spatial expression of vcan is essential for the development of various tissues such as the heart [24] or the joint interzone [36], or during interdigital web regression [37]. In these contexts, vcan plays two major roles in ECM architecture: (1) through its interaction with HA and chondroitin sulfate chains, it regulates matrix hydration and density, supporting the dynamic shaping of developing structures [38]; and (2) its cleavage by metalloproteases disrupts these interactions and promotes the emergence of specialized ECM fibers leading to matrix compaction [22] [39] [40]. Such remodeling events are essential for the structural maturation of tissues. For example, during heart development, the hydrated and loose matrix of the cardiac jelly is remodeled through vcan cleavage by ADAMTS1, a critical process for cardiac cushion morphogenesis [24]. A similar mechanism may occur during podocyte layer formation, where vcan is initially expressed during early assembly stages, but should be subsequently degraded through proteolytic activity to allow the maturation and functional organization of the podocyte layer. Interestingly, *adamts1* is specifically expressed in proximal region of the pronephros at stage 32 and may represent a good candidate involved in this remodeling process (S5 Fig). Notably, β1-integrin enrichment in the developing podocyte layer by stage 32, could be a consequence of this remodeling leading to pocket protrusion. Nevertheless, in absence of vcan, podocytes are present and correctly differentiated, as evidenced by *nphs2* expression. In mice, α3β1 integrin heterodimers anchor podocytes to the GBM, and β1-integrin deletion disrupts this adhesion, leading to podocyte loss [15] [16]. Altogether, these findings suggest that in the absence of vcan, podocytes fail to organize properly, and integrins may be unable to establish effective interactions between podocytes and the GBM.

It will be interesting in a future investigation to study the regulatory mechanisms governing *vcan* expression and its potential cleavage by ADAMTS proteases to fully elucidating its role in kidney morphogenesis.

## Acknowledgments

We thank S. Authier, E.Manzoni and A. Mannioui for excellent technical assistance in the maintenance of the *Xenopus* animal facility. We thank F. Lam and C. Chaumeton from the IBPS imaging platform.

## Supporting information captions

**S1 Fig. Vcan loss-of-function**

(A) Sequence of the morpholinos vcan-Mo1 and vcan-Mo2. The initiation codon ATG is highlighted in red and Mo sequences in grey. (B-J) Immunodetection of vcan on transverse sections of stages 32 embryos injected with St-Mo (B-D), vcan-Mo1 (E-G) or vcan-Mo2 (H-J). Panels (C, D), (F, G), and (I, J) are magnification of panels (B), (E) and (H) respectively. Scale bars: 280 μm for section and 93 μm for sections magnification.

**S2 Fig. Expression of the nephrostomes marker *odf3* in vcan-Mo1 depleted embryos at stage 35** WISH analysis of the nephrostomes gene marker *odf3* at stage 35. Representative embryos are shown.

*Odf3* labeling ratio between injected side and uninjected side appears similar, st-Mo= 48%, n=48 and vcan-Mo1= 50%, n=54. Scale bar: 850 μm.

**S3 Fig. Expression of the podocyte layer marker *nphs2* in vcan-Mo2 depleted embryos at stage 35** WISH analysis of the glomus gene marker *nphs2* at stage 35. Magnification of the pronephric region is highlighted by the dotted squares. (C, F) are transverse sections of representative St-Mo and vcan-Mo2 injected embryos with *nphs2* WISH. (Note: sections were not made from the whole mount specimen shown). Higher magnification of the tubule areas (indicated by dotted squares in C and F) are shown to the right of each image. Scale bars: 1.7 mm for whole embryo, 280 μm for embryo magnification, 280 μm for section and 93 μm for section magnification.

**S4 Fig. Expression of fibronectin and laminin in vcan depleted embryo at stage 32**

(A-L) Transverse sections of stage 32 embryo. Co-immunostaining of fibronectin (A-C) and laminin (D-F) in st-Mo injected embryos, and in vcan-Mo1 injected embryos (G-I for fibronectin; J-L for laminin). White arrows indicate the podocytes layer. Panels (B, C), (E, F) (H, I) and (K, L) are magnification of the pronephric areas indicated by dotted squares in panels (A), (D), (G) and (J), respectively. Scale bars: 280 μm for section and for sections magnification.

**S5 Fig. ADAMTS1 is expressed in the proximal pronephric part at stage 32**

ADAMTS1 expression *in X. laevis* embryos by WISH. The embryos are shown in lateral view with a magnification of the pronephric region indicated by the dotted squares. Scale bars: 1.7mm for whole embryo and 0.3 mm for magnification.

## List of abbreviations

(WISH): whole mount *in situ* hybridization
(ECM): extracellular matrix
(GBM): glomerular basement membrane

